# Immunization with recombinant enolase of *Sporothrix* spp (rSsEno) confers effective protection against sporotrichosis in mice

**DOI:** 10.1101/541029

**Authors:** Deivys Leandro Portuondo Fuentes, Paulo Roberto Dores-Silva, Lucas Souza Ferreira, Damiana Téllez-Martínez, Caroline Maria Marcos, Maria Luiza de Aguiar Loesch, Fanny Guzmán Quimbayo, Júlio César Borges, Alexander Batista-Duharte, Iracilda Zeppone Carlos

## Abstract

In recent years, research has focused on the immunoreactive components of the *S. schenckii* cell wall that can be relevant targets for preventive and therapeutic vaccines against sporotrichosis, an emergent worldwide mycosis. In previous studies, we identified a 47-kDa enolase as an immunodominant antigen in mice vaccinated with purified fungal wall proteins and adjuvants. In this study, the immunolocalization of this immunogen in the cell wall of *S. schenckii* and *S. brasiliensis* is shown for the first time. In addition, a recombinant enolase of *Sporothrix* spp (rSsEno) was studied with the adjuvant Montanide Pet-GelA (PGA) as a vaccine candidate. The rSsEno was produced with high purity. In addition, mice immunized with rSsEno plus PGA showed increased antibody titers against enolase and increased median survival time comparedto nonimmunized or rSsEno-immunized mice. Enolase immunization induced a predominant T-helper-1 (Th1) cytokine pattern in splenic cells after in vitro stimulation with rSsEno. Elevated production of interferon-γ (IFN-γ) and interleukin-2 (IL-2) was observed with other cytokines involved in the innate immune defense, such as TNF-alpha, IL-6, and IL-4, which are necessary for antibody production. These results suggest that we should continue testing this antigen as a potential vaccine candidate against sporotrichosis.

## Introduction

Sporotrichosis is a subcutaneous mycosis of subacute or chronic evolution caused by traumatic inoculation or the inhalation of spores of different species of the *Sporothrix* genus affecting both humans and animals^1^. The disease has a universal geographical distribution, although it is endemic in Latin America, including in Peru, México, Colombia, Guatemala and, especially, Brazil, where in the last 20 years, it became an important zoonosis, with the cat being the main source of transmission^2–4^. Species of the Sporothrix genus are thermodymorphic fungi with a saprophytic life at 25 °C and a filamentous form. The parasitic form at 35-37 °C is a yeast^1,5^. The human infection is acquired in two ways: traumatic inoculation through the skin with materials contaminated with *Sporothrix* spp or inhalation. Zoonotic transmission principally occurs from cats to humans^6^.

The genus Sporothrix is currently classified into two clades: i) the clinical clade, which includes *S. brasiliensis, S. globosa, S. luriei* and *S. schenckii* sensu stricto and ii) the environmental clade, composed mainly of species less pathogenic to man and animals, such as *S. mexicana, S. pallida* and *S. chilensis^7,8^.* Brazil is the only country that has reported all species of the clinical clade, and *S. brasiliensis* is the most virulent species^9,10^. This species is also the most prevalent during zoonotic transmission through deep scratches and bites from infected cats^8^. In this country, though sporotrichosis has been reported in most states, the disease is a neglected disease, particularly in the state of Rio de Janeiro (JR), where the largest number of cases has been reported, representing a serious public health problem^3^. The Oswaldo Cruz Foundation (Fiocruz), Rio de Janeiro, a referral center for the diagnosis and treatment of this mycosis, diagnosed over 4000 human and feline sporotrichosis cases between 1998 and 2012^11^. More recently, according to data from the epidemiological bulletin of 001/2018 of the sanitary vigilance service of RJ, during 2015 to 2018 (May), 3510 new cases were confirmed^12^, which shows a progressive increase in the incidence and prevalence of this mycosis.

Sporotrichosis is usuallycontrolled through the combined use of itraconazole/potassium iodide or terbinafine in immunocompetent patients who exhibit the less severe clinical forms of the disease (lymphocutaneous and fixed cutaneous lesions)^13,14^. However, in immunocompromised patients with neoplastic diseases, transplantation or AIDS, the conventional treatment with classical antifungals is generally ineffective ^15,16^. The lack of a veterinary and/or human vaccine against this disease has awakened interest in the identification of *S. schenckii* cell wall immunoreactive components involved in fungal pathogenesis^17^ and the induction of the immune response^18^ that can be used for immunoprophylaxis and immunotherapy against sporotrichosis.

In previous studies, our group showed that sera obtained from mice immunized with an *S. schenckii*-cell wall protein (CWP) formulated with the adjuvant aluminum hydroxide (HA) showed reactivity against two proteins, one of 71 kDa and another of 47 kDa. The latter was functionally identified as enolase and predicted to be an adhesin by the Fungal RV database^19^. These immune sera showed opsonizing properties, enhancing the phagocytosis of *S. schenckii,* and they inhibited the fungal adhesion to fibroblasts *in vitro.* Passive transfer of immune serum to nonimmunized mice conferred protection against challenges with the fungus. These findings indicated the induction of protective immunity from the vaccine formulation against experimental sporotrichosis and the potential use of both antigens for an antifungal vaccine. More recently, we showed that serum from mice vaccinated with AH-adsorbed CWPs, and serum obtained from mice immunized with the same antigenic source but formulated with Montanide™ Gel Pet A adjuvant (PGA), reacted with the *S. brasiliensis* yeast cell wall^20^. Such cross-reactivity, as well as the fact that both formulations confer protection in mice challenged either with *S. schenckii or S. brasileinsis,* suggested the existence of shared immunodominant antigens that could prove beneficial for the simultaneous protection against these species, which are the more virulent of the genus *Sporothrix.*

Enolase (2-phospho-D-glycerate hydrolase, EC 4.2.1.11) is a metalloenzyme that requires the metal ion magnesium (Mg^2^+) to catalyze the dehydration of 2-phosphoglycerate (2-PG) to phosphoenolpyruvate (PEP), a product that is used to produce energy (ATP) in eukaryotic and prokaryotic cells^21^. In mammals, there are at least 4 subunits of enolase: α-enolase (eno1), expressed in almost all tissues; β-enolase (eno3), predominantly expressed in adult skeletal muscle; γ-enolase (eno2), found in neurons and neuroendocrine tissues^22^; and eno4, expressed in human and mouse sperm ^23^. Enolase has been identified on the cell surface of *C. albicans^24^, Plasmodium falciparum^25^, Ascaris suum^26^, Streptococcus sobrinus^27^, S. suis serotipo* II^28^, S. *iniae^29^, Plasmodium spp^30^* and *Clonorchis sinensis*^31^. In addition, the immunogenicity and protective properties of anti-enolase immune response have been reported for diverse pathogens^24,27,32^.

In this study, we reported for the first time an enolase enzyme in the cell wall of *Sporothrix* spp that is an immunogenic fungal cell wall component. We also showed the enzyme’s recognition by the serum of cats with sporotrichosis. A recombinant *S. schenckii* enolase was obtained, and it was used to prepare an enolase-based vaccine formulated with PGA adjuvant against *S. brasiliensis* in mice. The immunogenicity and protective capacity of enolase was observed in an anti-sporothrix prophylactic vaccine candidate in mice.

## Results

### Production, purification, and characterization of rSsEno

Figure 1A shows that rSsEno expressed in the IPTG-induced pET28a::SsEno-transformed-BL21 cells was produced both in the pellet, as well as in the soluble fraction of the lysed cells (Fig. 1, lane 4 and lane 5, respectively). Based on this result, the rSsEno containing the soluble fraction was purified by Ni^2+^-affinity (results not shown) and preparative size exclusion chromatography (Fig. 1, lane 6), respectively, resulting in an apparent purity of 95% on SDS-PAGE with Coomassie blue staining. rSsEno had the expected molecular weight of 47 kDa.

**Fig. 1.**
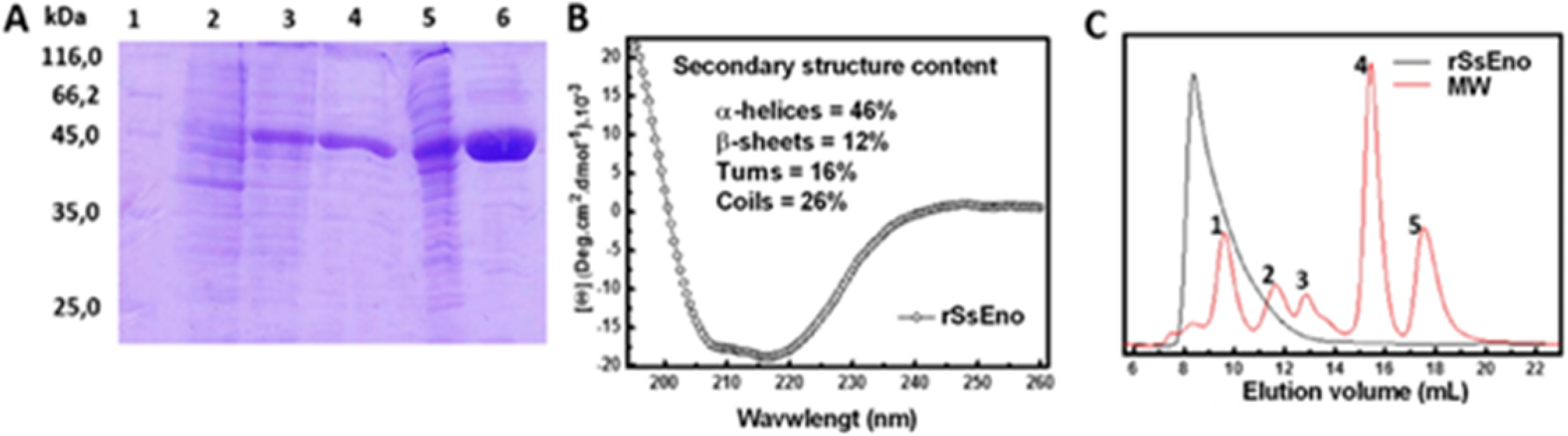
SDS-PAGE and structure analysis of rSsEno expressed in *E. coli* BL21. The recombinant plasmid pET28a::SsEno-transformed *E. coli* BL21 cells were induced in the presence of 0,2 mM IPTG for 4h at 30 °C. The cells were lysed by sonication, and the supernatant with the recombinant protein was purified by affinity and molecular exclusion chromatography, respectively. All the samples were analyzed by SDS-PAGE 12%, and the protein was stained with Coomassie Blue R250 in the gel. (A) Expression and purification of rSsEno. Molecular mass markers in kDa (1), non-IPTG-induced pET28a::SsEno-transformed-BL21 cells lysate (2), IPTG-induced pET28a::SsEno-transformed-BL21 cells lysate (3); pellet of IPTG-induced pET28a::SsEno-transformed-BL21 cells (4), supernatant of lysed IPTG-induced pET28a::SsEno-transformed-BL21 cells (5), rSsEno purified by Ni^2+^ affinity chromatography (6) and size exclusion chromatography (7). (B) The CD spectrum shows that rSsEno was obtained mainly with a secondary structure composed by α-helices. (C) Analytical size exclusion chromatography performed for rSsEno. The MW standard protein mix elution pattern is represented by the red line: 1) Apoferritin (480 kDa); 2) γ-Globulin (160 kDa); 3) BSA (67 kDa); 4) carbonic anhydrase (29 kDa); 5) Cytochrome C (12 kDa).

The rSsEno far UV-CD spectrum shows a positive band at ~195 nm and negative bands at ~209 and 218 nm, which indicate the presence of a folded structure into α-helices and β-sheets (Fig. 1B). Deconvolution analysis of the rSsEno far UV CD spectra revealed that the secondary structure of this protein contains higher α-helices (46%) and a smaller number of β-sheets (12%). Analytical SEC analysis showed that rSsEno elutes in the column void with a tail after the main peak; this suggests that the recombinant protein was organized by various oligomeric forms (Fig. 1 C).

### Sequence Alignment of the *S. schenckii* enolase

The sequence alignment analysis among the *S. schenckii-, F. catus-* and *H. sapiens*-enolase revealed an expected result; the enolase from humans and cats showed a degree of identify of 95% (Fig. 2). However, both enolases showed an identity of 62% with the enolase of *S. schenckii.*

**Fig. 2.**
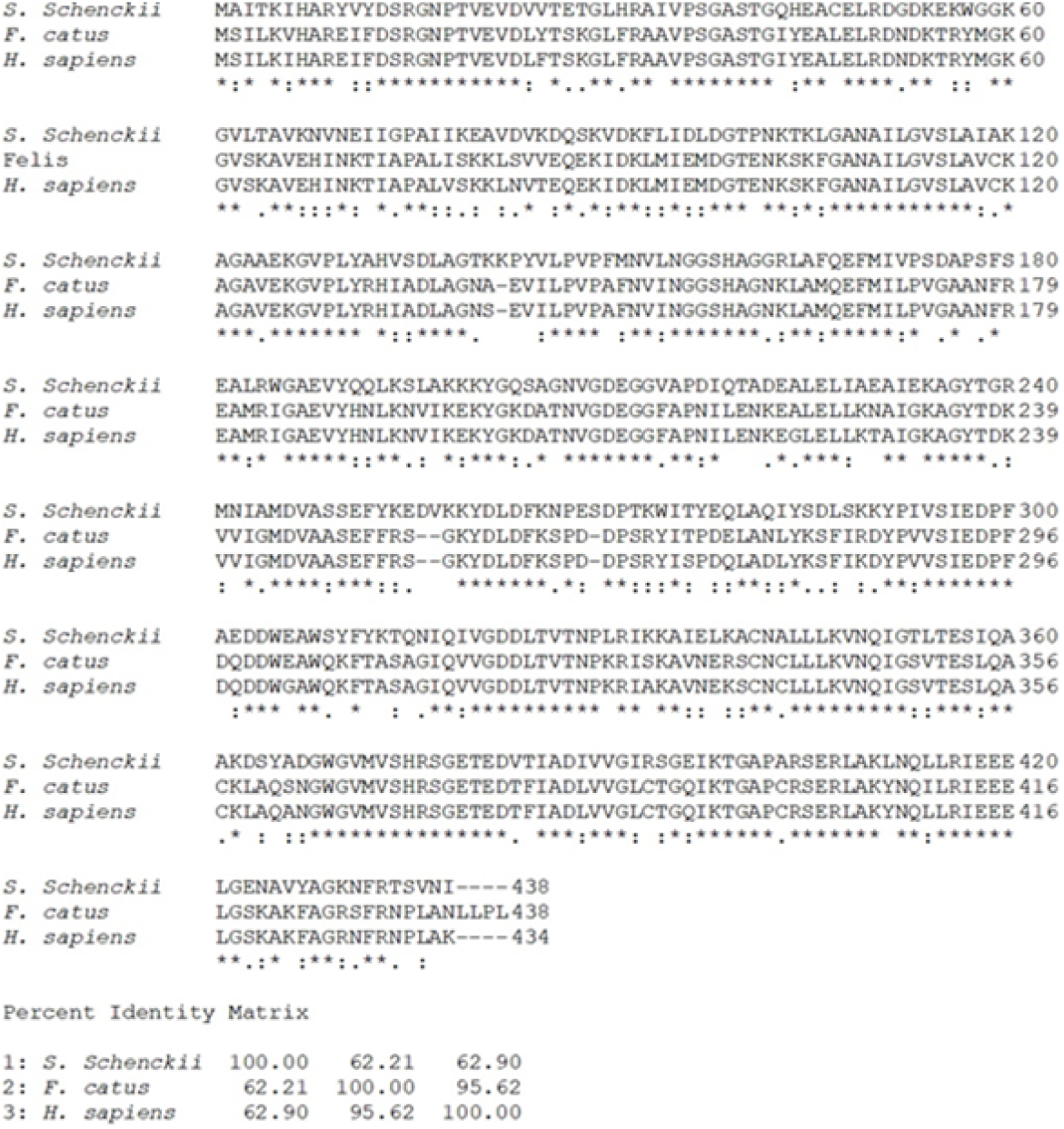
Multiple sequence alignments of *S. schenckii.* The deduced amino acid sequence of *S. schenckii* (ERS97971.1), *Felis catus* (M3WCP0_FELCA) and *Homo sapiens* (P06733) were aligned by the Clustal Omega server. The conserved amino acids in all sequences are labeled with asterisks; the conservative and semi-conservative substitutions are labeled with two and one points, respectively. The percentage of amino acid sequence identity between all enolases is indicated.

### Specificity of the anti-rSsEno serum

The specificity of the antibodies raised in rSsEno immunized mice was examined by immunoblotting against recombinant enolase or CWP isolated from Ss16345. As shown in Figure 3 (A), the anti-rSsEno serum reacted with the recombinant protein and against a single reactive band present in Ss16345CWP with the expected 47 kDa molecular mass, corresponding to the native enolase^19^. Interestingly, the serum obtained from infected cats with sporotrichosis confirmed a specific high reactivity against the recombinant protein (Fig. 3B and C), indicating that, during natural infection, the fungal enolase can induce anti-enolase antibodies. Sera from uninfected control cats exhibited no immunoreactivity with the rSsEno (Fig. 3B)

**Fig. 3.**
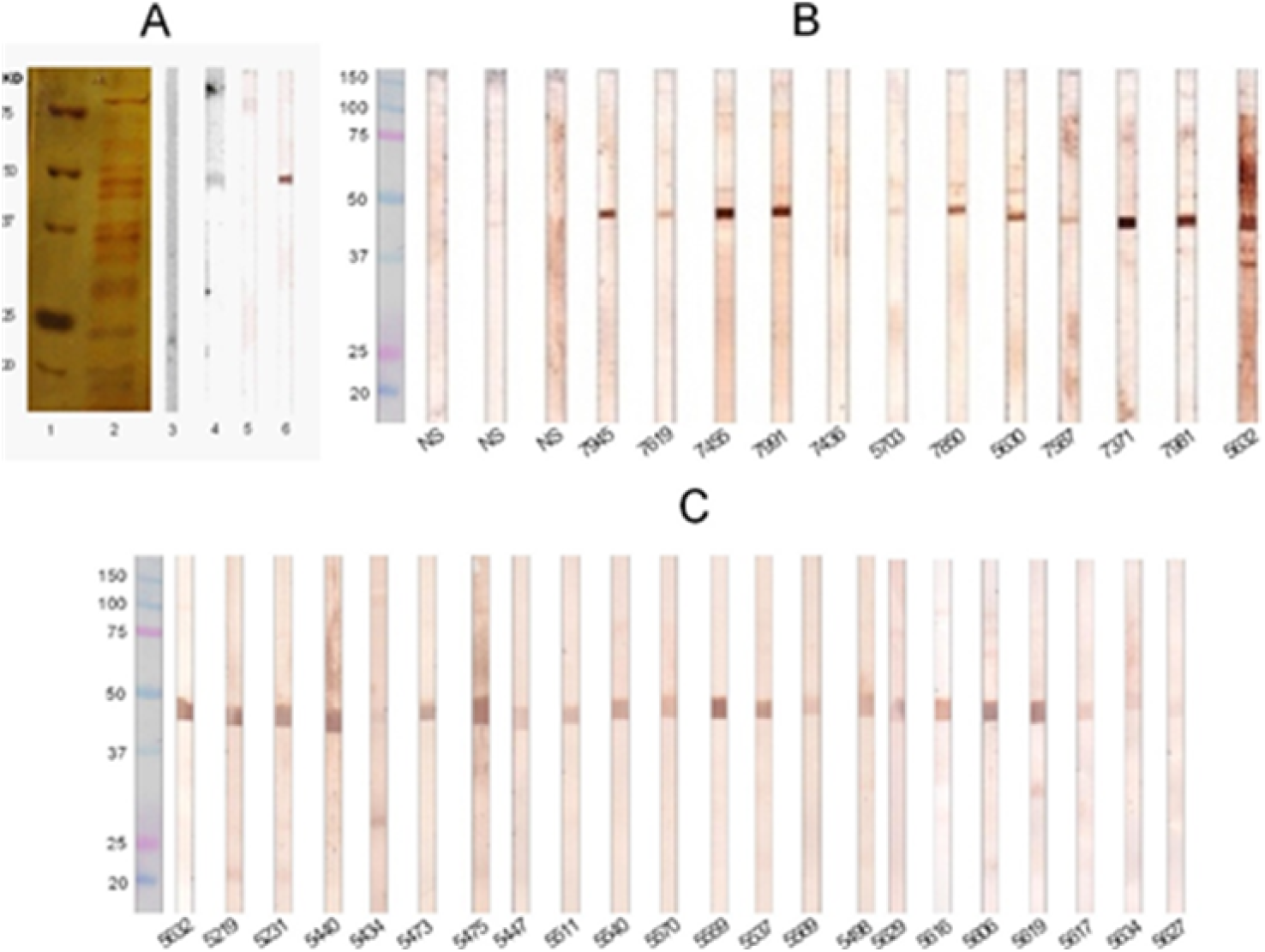
Western blot analysis showing the specificity of the anti-rSsEno sera and the reactivity of the sera from cats with sporothricosis against rSsEno. Samples Ss16345-CWP and rSsEno were tested by 12% SDS-PAGE under nonreducing conditions and after immobilization on a nitrocellulose membrane. The strips were incubated at 37 °C for 1 h with anti-rSsEno serum or naïve mouse serum, and the immunoblots were visualized by adding 3,3′-diaminobenzidine substrates after being treated with goat anti-mouse IgG-HRP. Panel A, column 1: molecular weight marker; column 2: Ss16345-CWP resolved by SDS-PAGE 12%. column 3 and 4: nitrocellulose strips containing the Ss16345-CWP treated with NS and anti-rSsEno serum, respectively. Column 5 and 6: nitrocellulose strips containing rSsEno treated with naïve mice-serum and anti-rSsEno serum, respectively. Panel B, strips containing rSsEno were incubated with sera from cats with or without sporotrichosis (NS) and immunoblots were incubated with goat anti-feline IgG-HRP. Each cat serum is identified by the admission number of the Laboratory of Clinical Research in Dermatozoonoses in Domestic Animals of the National Institute of Infectology Evandro Chagas (FIOCRUZ). The resulting blots were cropped to show the bands of interest and received equal exposure levels.

### Enolase is present in the cell wall of *S. schenckii* spp

After confirming their specificity, the anti-rSsEno serum was used to detect enolase in the Ss16345, Ss1099-18, Ss250 and Ss256-cell wall. Figure 4 (A, C, E and G) shows an intense and significant (p<0.05) median fluorescence intensity (MFI) in yeasts treated with the anti-rSsEno serum compared to yeast treated with serum from nonimmunized mice (NIS), evidencing enolase on the cell surface of these strains. The MFI was higher in the cat isolate (Fig. 4F and H) compared to Ss16345 (Fig. 4B) and Ss1099-18 (Fig. 4D), suggesting that this protein is expressed more on the cell wall of *S. brasiliensis,* the more virulent species. The presence of enolase on the cell surface of the studied fungi was also confirmed by transmission microscopy using the immunogold stain. Figure 4 showed that enolase appears distributed along the cell wall of Ss16345, Ss1099-18, Ss250 and Ss256, which might facilitate its recognition by the host’s immune system, although it also appears, as expected, in the cellular cytoplasm of these species, since its classical function is to catalyze the reversible conversion of 2-phosphoglycerate to phosphoenolpyruvate^22,33^.

**Fig. 4.**
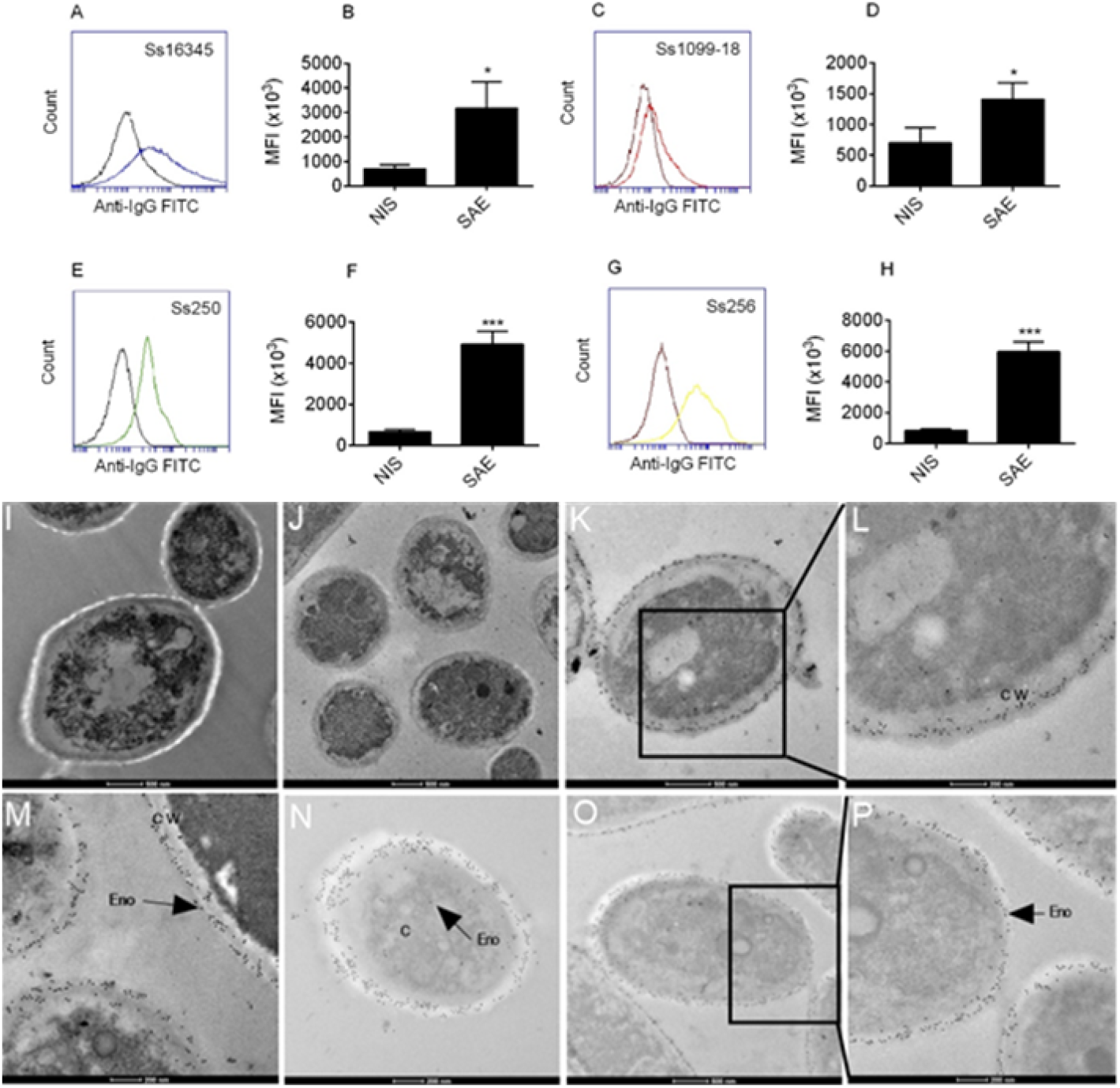
Demonstration of the enolase on the cell surface of *Sporothrix sp-yeasts* by flow cytometry and electron microscopy. A Ss16345, Ss1099-18, Ss250 and Ss256 yeasts suspension was previously incubated with anti-rSsEno serum (SAE) or serum from nonimmunized mice (NIS) for 1 h at 37 °C. After washing, the cells were exposed to FITC rabbit anti-mouse IgG and examined using a flow cytometer. (A, C, E and G) Representative histograms from one of three independent experiments for the indicated *Sporothrix* spp yeasts treated with NIS (black line) or SAE (blue, red, green or yellow line). Bar graphs show the median fluorescence intensity (MFI) for Ss16345 (B), Ss1099-18 (D), Ss250 (F) and Ss256 (H). The results are presented as the mean ± SD of three independent experiments, and statistical significance was determined by a’s student paired *t* test. *, *P* < 0.05; *** *P* < 0.001. Ultrathin sections of each fungus were incubated overnight with anti-anti-rSsEno (SAE) serum or serum from nonimmunized mice (NIS) following treatment overnight with the Au-conjugated secondary antibody, rabbit IgG, at 4 °C. Grids were observed with a Jeol 1011 transmission electron microscope after being stained with uranyl acetate and lead citrate. (I) shows Ss16345-yeasts. (J) shows Ss16345-yeasts treated with NIS. (K and L) show the enolase (Eno) on the cell wall (CW) or cytoplasm (C) of the Ss16345-yeasts treated with SAE. (M) shows the Eno on CW of the Ss1099-18-yeasts treated with SAE. (N) shows Eno in the cytoplasm or CW of the Ss250-yeasts treated with SAE, respectively. (O and P) representative images showing Eno on the CW of the Ss256-yeasts treated with SAE.

### Antibody response

To assess the immunogenic potential of *S. schenckii*-enolase, sera from the experimental group obtained seven days after the last boost was subjected to ELISA using rSsEno as an antigen. Our results showed that animals immunized only with enolase stimulated high IgG specific antibody production (Fig. 5A) compared to the PBS group. However, as expected, the specific antibody production was significantly higher (p <0.05) when enolase was formulated with the PGA adjuvant. We also determined the rSsEno-specific IgG1, IgG2a and IgG3 antibodies induced by each formulation. Mice immunized with rSsEno100 and PGA+rSsEno100 induced higher IgG1 and IgG3 antibody levels against rSsEno compared to the PBS control group, but the level of both subclasses was higher in the mice immunized with the PGA-adjuvanted formulation (Fig. 5B and C). The PGA+rSsEno100 formulation was the only formulation that induced the production of IgG2a (Fig. 5D).

**Fig. 5.**
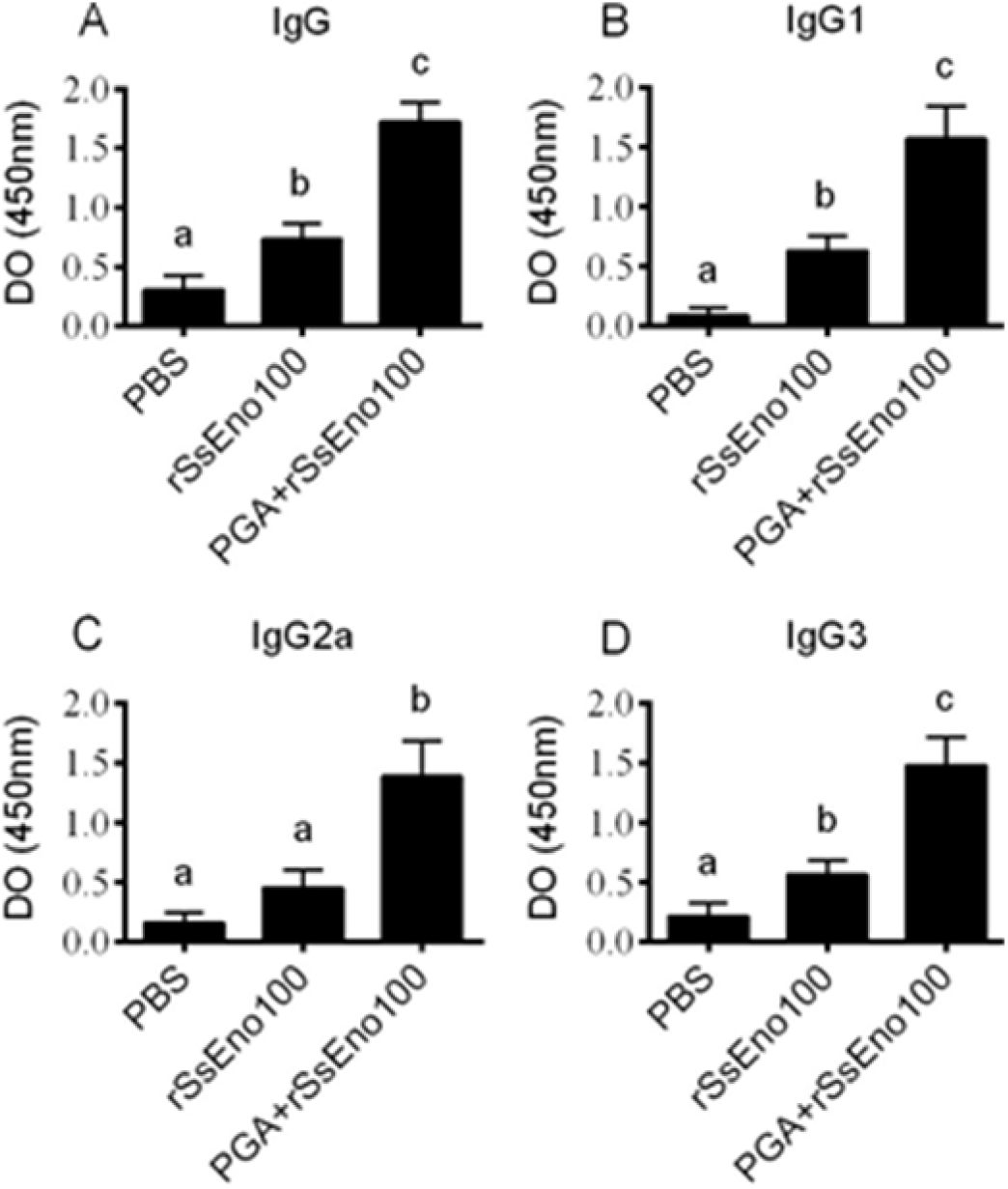
Immunization with rSsEno with or without PGA conjugation enhanced the antibody response. BALB/c mice were s.c. immunized three times with rSsEno100, PGA+ rSsEno100 or PBS as a negative control. Sera collected seven days after the last boost was used to determine antigen-specific IgG (A), IgG1 (B) and IgG2a (C) titers by ELISA. The results are presented as the mean ± SD of 5 mice from one of three independent experiments, and statistical significance was determined by one-way ANOVA using Tukey’s multiple comparisons test and a 95% confidence interval. Different letters represent significant differences (p<0.05) between treatments.

### Cytokine profile analysis

The effect of anti-enolase vaccination on the pattern of cytokines was evaluated in the supernatant of splenocyte cultures from nonimmunized and immunized mice after in vitro stimulation with rSSEno100. A higher production of IL-2 and IFN-ɣ from the Th1 profile, IL-4 and IL-6, which are involved in the production of antibodies, and TNF-α, which is released during the innate immune response (also with IL-6) in mice vaccinated with PGA+rSsEno100, was observed (Fig. 6). All of these cytokines are involved in defense against *S. schenckii,* which is additional evidence of protective immunogenicity induced by the vaccine formulation.

**Fig. 6.**
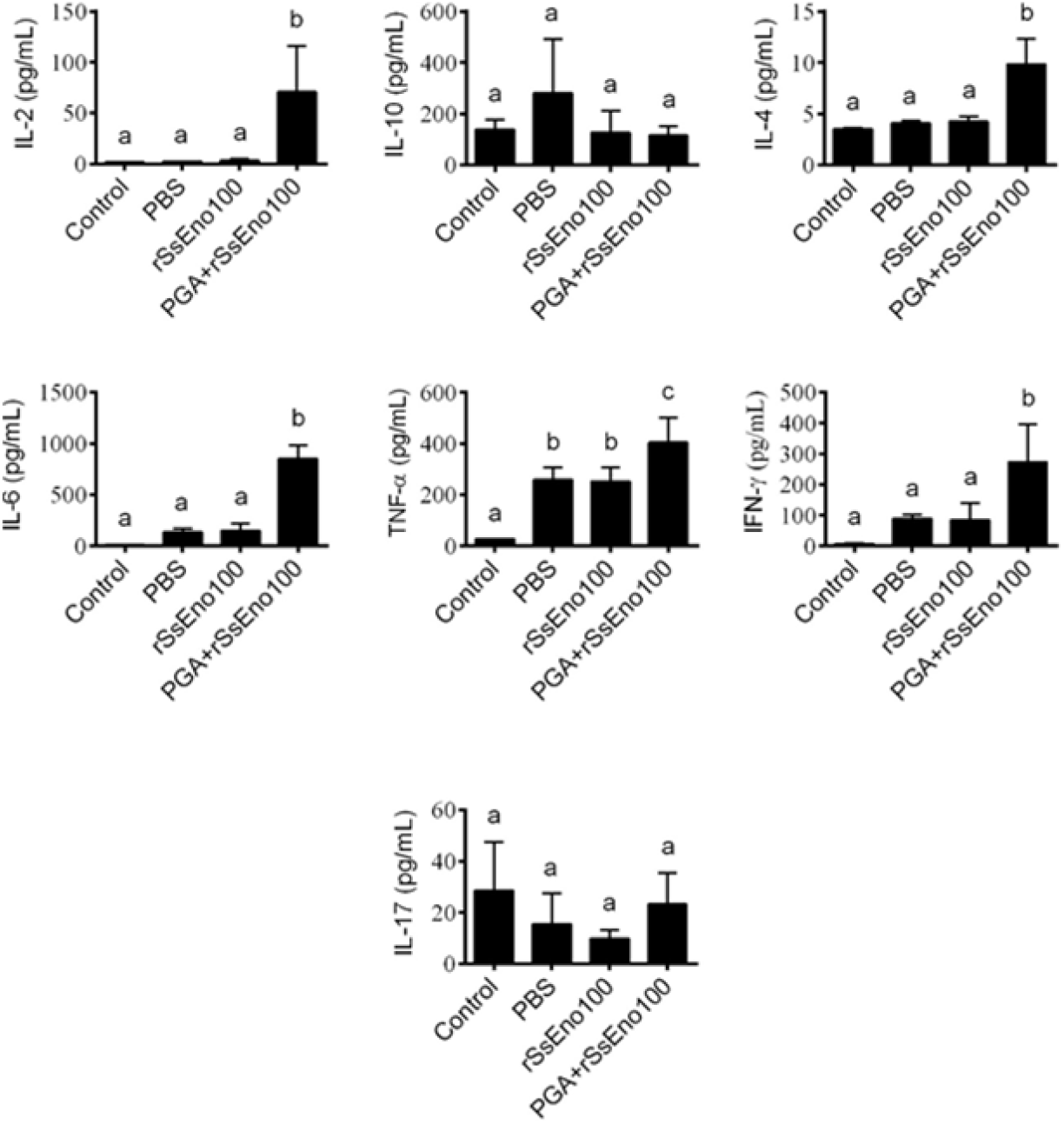
Vaccinated mice with rSsEno100 and PGA+rSsEno100 showed differences in Th1, Th2 and Th17 cytokine profiles. BALB/c mice were s.c. immunized three times with rSsEno100, PGA+rSsEno100 or PBS as a negative control. Total splenocytes of each animal were obtaining seven days after the last immunization and stimulated in vitro with rSsEno. After 24 h of incubation, supernatant-accumulated cytokines (IL-2, IL-4, IL-6, IL17A, IFN-γ, TNF and IL-10) were measured by cytokine cytometric bead array kit ELISA. The results are presented as the mean ± SD of 5 mice from one of three independent experiments, and statistical significance was determined by one-way ANOVA using Tukey’s multiple comparisons test and a 95% confidence interval. Different letters represent significant differences (p–0.05) between treatments.

### Challenge studies

To test whether rSsEno in the formulation with PGA adjuvant protects against systemic sporotrichosis in mice, seven days after booster immunization, mice from each group were challenged intravenously with 10^5^ *S. brasiliensis* 250 yeasts, a highly virulent specie. The mortality of nonimmunized mice was of 100% before 40 days postinfection, while the rSsEno-immunized mice showed over 50% survival, and those immunized with the PGA-adjuv anted formulation exhibited the highest percentage of survival (over 90%) at the end of the experiment (45 days postinfection) (Fig. 7). These results clearly show that enolase may have a potential use for vaccination against sporotrichosis.

**Fig. 7.**
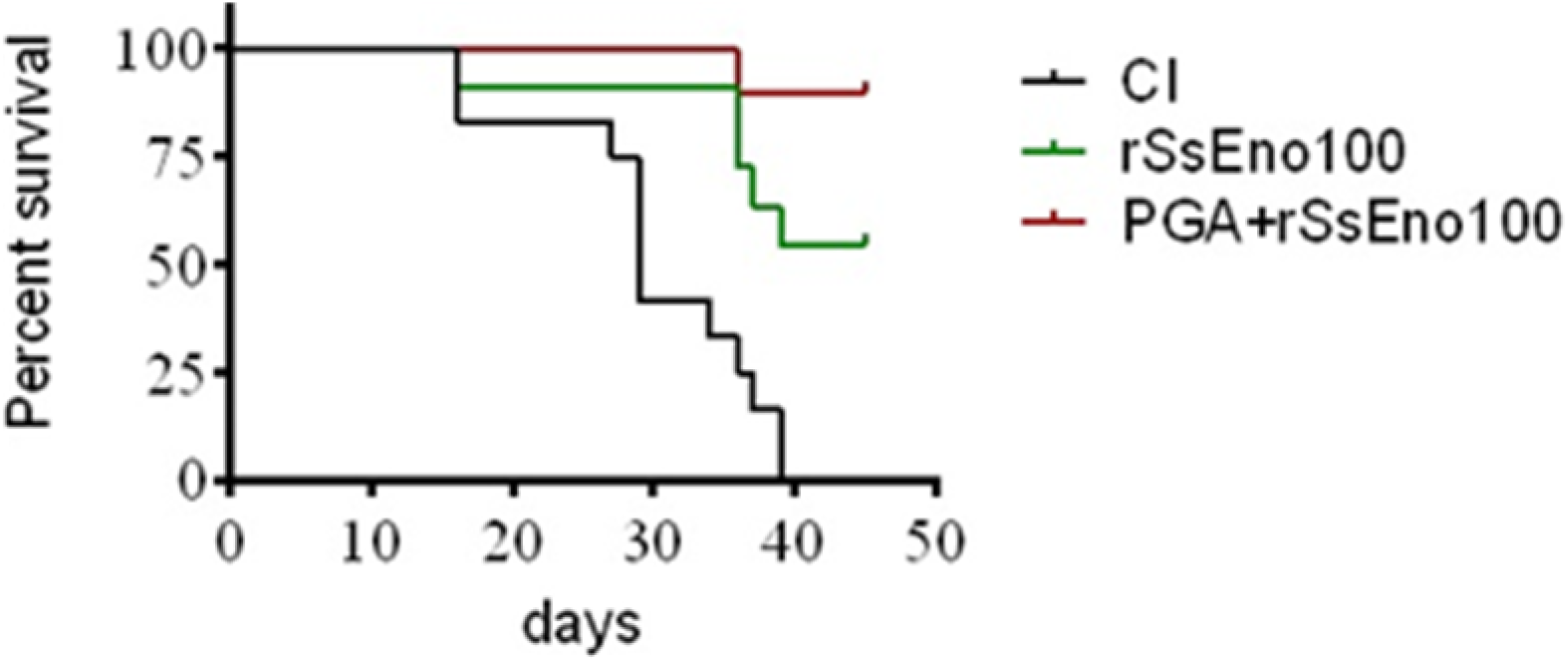
Vaccination of mice with rSsEno with or without PGA adjuvant conferred protection against infection with *S. brasiliensis* (Ss250). BALB/c mice were immunized (s.c.) three times with the indicated formulations. One week after the last boost, mice were challenged intravenously with 1×10^5^ Ss250 yeast-form cells. The survival of the mouse groups was monitored daily for 45 days postchallenge (n=10 in all groups).

## Discussion

In the last two decades, sporotrichosis has been a hyperendemic zoonosis transmitted preferentially by the domestic cat, especially in Brazil. The high incidence of sporotrichiosis together with the ineffectiveness of treatment, especially in immunocompromised individuals, has reinforced the need to identify antigenic targets on the cell surface of species of clinical interest of the genus Sporothrix for immunological prevention and therapeutic intervention ^17,34^

In this study, the enolase of *S. schenckii* was obtained by recombination with *E. coli,* and it was purified and characterized. Our results showed that rSsEno was produced successfully with the expected molecular weight of 47 kDa and with the secondary structure of a type α-helix at 46% and a β-sheet at 12%. These values are close to values of 43% for the α-helix and 15% for the β-sheet from *Saccharomyces cerevisiae* enolase^35^, suggesting that both enolases have similar structures. The analyses of the oligomeric state of rSsEno by gel filtration revealed an unexpected result. We expected that this protein would be assembled in the form of dimers, as reported for yeast enolase ^36–38^. However, our results indicate that the rSsEno was produced with a molecular weight of 480 kDa, ten times greater than the 47 kDa monomeric unit of this protein.

Enolase has been described as a moonlighting protein that exhibits multiple nonglycolytic functions, probably because of its different multimeric structures^32^. Ehinger *et al.* ^39^reported that a-enolase of *Streptococcus pneumonia* forms an octamer in solution and that due to its binding to human plasminogen, it probably resides on the cellular surface of this pathogen and can be involved in virulence. Wu *et al*. ^40^ also reported that *Staphylococcus aureus* recombinant enolase is organized in dimers and octamers and that the latter probably exist *in vivo* since it showed enzymatic activity *in vitro.* Whether the complex oligomeric state of rSsEno in solution is the same as its native form in *S. schenckii,* and its functional role *in vivo,* is a subject for future studies.

The reactivity of sera from cats with sporotrichosis against rSsEno and the lack of reactivity with sera from uninfected control cats evidenced the antigenic role and probable immunogenicity of the *S. schenckii* enolase during the infectious process in these animals. This result, in addition to the percentage of identity (62%), suggests that *S. schenckii* enolase contains conserved regions distinct to the enolase present in both hosts (cat and human) of this pathogen. Therefore, enolase can be an antigenic target for vaccine and/or therapeutic strategies for protection against sporotrichosis in cat.

Different studies have shown that enolase on the cell surface of bacteria, fungus and parasites acts as a virulence factor that facilitates the colonization and dissemination of these pathogens in the host^25,38,41^. In this study, we show for the first time that enolase is present on the cellular surface of *S. schenckii* and *S. brasiliensis* species, and interestingly, this expression was higher on yeast cell walls from *S. brasiliensis,* suggesting that the level of enolase expression on the cell surface of species of the genus Sporothrix can be related to the invasiveness and virulence of these pathogens in the host. In this way, Roth *et al.* ^42^ showed that the level of expression of enolase is 15-fold higher in red blood cells infected with *P. falciparum* compared to uninfected cells. More recently, Marcos *et al.*^43^ observed a considerable increase of this protein in the cell wall of *Paracocidiodes brasiliensis* when the fungus was cultivated in BHI medium enriched with sheep blood or during fungal infection in mice, suggesting a role for enolase as a virulence factor of these fungi in host cells.

Several studies have shown that the IgG antibody response^44,45^ and especially IgG1^46,47^, IgG2a and IgG3 isotypes^19,20^ against the *S. schenckii* and *S. brasiliensis* cell wall proteins is associated with protection against progressive infection. Our results showed that rSsEno100 and PGA+rSsEno100 stimulated a Th2 (IgG1 and IgG3) and Th1/Th2 (IgG1, IgG2a and IgG3) immune response, respectively.

The generation of a Th1 and Th17 response is necessary for protective immunity against *Staphylococcus aureus* and *C. albicans*^48^. Ferreira *et al.^49^* demonstrated in a model of *S. schenckii* infection in BALB/c mice that the Th1 and Th17 immune response were able to control the infection. Recently, our group reported a similar result in a model of C57BL6 mice subcutaneously infected with either *S. schenckii* or *S. brasiliensis*. However, the higher virulence of *S. brasiliensis* caused a long-lasting infection associated with severe tissue lesions that stimulated a regulatoryT cell (Tregs) response with deleterious effects on the Th1 and Th1/Th17 response, although a compensatory Th17 response was induced^50^. We also demonstrated in an immunoprophylaxis study in BALB/c mice that either aluminum hydroxide adjuvant or PGA, both formulated with the Ss16345-WCP containing the immunoreactive enolase, induced a Th1, Th2 and Th17 profile, in addition to high stimulation of specific antibodies that conferred protection in these animals after challenge with Ss16345 or Ss250^20^.

To verify whether rSsEno could be used as an antigenic target for a sporotrichosis vaccine, we performed a survival study in immunized mice after intravenous infection with the highly virulent strain Ss250. The survival above 90% seen in mice immunized with PGA+rSsEno100 is strong evidence of the protective capacity of our vaccine candidate. In addition, the Th1/Th2, and not the Th17 cytokine profile, observed ex vivo in PGA+rSsEno100-immunized mice played a significant role in vivo in favoring protection, since rSsEno100-immunized mice showed ex vivo that the stimulation of Th2 cytokines alone led to decreased survival (~ 52%) postchallenge. Li *et al.*^24^ showed that a Th1 and Th2 immune response pattern induced by recombinant enolase of *C. albicans* emulsified with Freund’s adjuvant (AF) was enough to confer protection on C57BL/6 mice challenged with a lethal dose of *C. albicans* strains SC5314 and 3630. In addition, passive immune serum transfer, characterized by the prevalence of IgG2a- and IgG1-specific antigen isotypes, also demonstrated effective protection against both fungal *C. albicans* lineages, showing that antibodies against enolase could be useful to treat of candidiasis. Zhang *et al*.^51^ also showed that the enolase of *Streptococcus suis* serotype 2 plus AF formulation induced a mixed Th1 (IgG2a) and Th2 (IgG1) response that also conferred protection in challenged animals with two pathogenic strains of *S. suis.* This same immune response profile and protective efficacy were observed in mice immunized with the *Ascaris suum* enolase after infection with infective larvae of this parasite^52^.

In summary, for the first time, a recombinant form of *S. schenckii* (rSsEno) enolase was obtained and structurally characterized. The molecular mass of rSsEno determined by size exclusion chromatography was 480 kDa, which shows that this protein is organized as more than two monomeric units. This organization is different from the enolases from other fungi. The identification of enolase on the cell wall of *S. brasiliensis* and *S. schenckii* and its recognition by serum from cats affected with sporotrichosis are reported in this study. A vaccine formulation of rSsEno plus PGA adjuvant induced a Th1/Th2 response and high titers of specific antibodies that favored the protection to mice challenged with a highly virulent *S. brasiliensis* isolate. All these results show that the enolase of *Sporothrix* spp may be a vaccine antigen candidate for feline sporotrichosis prevention.

## Materials and methods

### Animals

For this study, male 5-7-week-old BALB/c mice were purchased from “Centro Multidisciplinar para Investigação Biológica na Área da Ciência de Animais de Laboratório” (CEMIB), Universidade de Campinas (UNICAMP), São Paulo, Brasil. Animals were housed in individually ventilated cages in an ambient controlled temperature and 12-h light/dark cycles. All animals were acclimatized to the conditions for 1 week before the experiments, and water and food was offered ad libitum. This study was carried out in strict accordance with the recommendations for the Guide for the Care and Use of Laboratory Animals of the National Institutes of Health, and the protocols were approved by the Ethics Committee for Animal Use in Research of Araraquara’s School of Pharmaceutical Sciences from UNESP (Protocol CEUA/FCF/CAR no. 57/2015).

### Microorganisms

The strains *S. schenckii* ATCC 16345 (Ss16345), *S. schenckii* 1099-18 (Ss1099-18), *S. brasiliensis* 250 (Ss250, GenBank: KC693883.1) and S. *brasiliensis* 256 (Ss256, KC693889.1), both *S. brasiliensis* strains isolated from feline sporotrichosis, and Ss16345 were kindly provided by the Oswaldo Cruz Foundation, Rio de Janeiro, Brazil. Ss1099-18 was provided by Dr. Celuta Sales Alviano at the Institute of Microbiology, Federal University of Rio de Janeiro (Brazil). Mycelial-to-yeast phase conversion was accomplished as previously described by Ferreira and collaborators^49^.

Expression and purification of recombinant *S. schenckii* Enolase (rSsEno)The gene that encodes *S. schenckii* enolase with 438 amino acids and a molecular mass of 47 kDa (Accession Code: ERS97971.1 of the GenBank database) was synthesized by Epoch Life Science Inc. between the *Nde I* and *Eco RI* restriction enzymes in fusion with a histidine tag at the N-terminus. It was subcloned into the pET28a plasmid and optimized for production in *E. coli* (pET28a::SsEno).

*Escherichia coli* DH5α was used as the cloning host for the propagation of pET28a::SsEno on lysogeny broth (LB) agar medium containing 30 μg/mL of kanamycin, and the authenticity of the cloning procedure was confirmed by sequencing. For recombinant protein expression, *E. coli* BL21 cells cotransformed with pET28a::SsEno were grown at 37 °C in LB medium containing 30 μg/mL of kanamycin until they reached an OD_600_ in the range of 0.5-0.7. The expression of rSsEno was induced by 0.2 mmol/L of isopropyl β-D-1-thiogalactopyranoside (IPTG) at 30°C for 4 h. The cells were separated by centrifugation for 20 min at 8000 rpm, and the pellet was resuspended in 20 mL buffer A (NaPO_4_ 20 mM, NaCl 500 mM and imidazole 20 mM, pH 7,4) containing 5 U of DNAse (Promega) and 30 ug/mL lysozyme (Sigma) for 30 min on ice. The cell homogenate was sonicated, filtrate and then centrifuged at 19,000 rpm for 20 min at 4 °C. The rSsEno-containing supernatant was filtered through a 0.45 μm nitrocellulose membrane (Millipore) and further subjected to Ni^2+^-affinity chromatography in buffer A. The rSsEno was then eluted in buffer B (NaPO4 20 mM, NaCl 500 mM and imidazole 500 mM, pH 7,4). After elution, the material obtained was subjected to size exclusion chromatography (SEC) with a Superdex 200 pg 16/60 column (GE Healthcare Life Sciences) in Tris-HCl 25 mM, NaCl 100 mM and β-mercaptoethanol 2 mM at pH 7.5, and the eluted protein was concentrated using the *Amicon^®^ Ultra 15 mL 3k* device (Millipore) after being dialyzed for 24 h at 4°C against phosphate buffer saline (PBS, pH 7,2-7,4). The rSsEno concentration was measured by the BCA assay (Pierce), and the efficacy of the expression and purification processes was assessed by 12% SDS-polyacrylamide gel electrophoresis (SDS-PAGE).

### Circular dichroism (CD)

Secondary structure analysis was performed by far-UV (195-260 nm) CD in a J-815 spectropolarimeter (Jasco Inc.) coupled to a Peltier PFD 425S for the temperature control system. rSsEno was tested in Tris-HCl buffer (pH 7.5), 100 mM NaCl and 2 mM β-mercaptoethanol, and the secondary structure content was estimated using the CDNN Deconvolution program^53^. In addition, the rSsEno oligomeric state was analyzed by analytical size exclusion chromatography on a Superdex 200 GL 10/30 column (GE Healthcare LifeSciences) coupled to a ÄKTA Prime Plus (GE Healthcare LifeSciences) and equilibrated with the same buffer described above.

### Extraction of Ss16345CWP

Extraction of the Ss16345CWPs was performed per Portuondo *et al*.^19^. Ss16345 yeast cells collected from logarithmically growing cultures were incubated with a protein extraction buffer containing 2 mM dithiothreitol, 1 mM phenylmethylsulfonyl fluoride, and 5 mM EDTA in This-HCl buffer for 2 h at 4°C under mild agitation. The Ss16345CWP-containing supernatant was collected, dialyzed against PBS, and then concentrated using the Amicon Ultra 15 MWCO concentrator (Millipore). The proteins were then precipitated by overnight incubation with 10% (w/v) trichloroacetic acid in acetone at 4°C, and the resulting pellets were washed in ice-cold acetone, dried in a SpeedVac^®^ and reconstituted PBS. The protein concentration was measured by the BCA assay (Pierce).

### SDS-PAGE, Western blot analysis

Samples containing 20 μg of protein Ss16345CWP and purified rSsEno (5 μg) were resolved on an SDS-PAGE 12% as described by Laemmli^54^. Two gels were stained with Coomassie brilliant blue R250, and the other gels were transferred to 0.45-μm-nitrocellulose membranes (GE Healthcare) using a mini Tank VEP-2 electroblotting system (Owl Separation Systems, Thermo Scientific) at 50 mM for 3 h. The membrane-cut strips were saturated with 5% dried skim milk in PBS for 4 h at 37°C, and the strips containing rSsEno were incubated overnight at room temperature (RT) with anti-rSsEno serum (obtained from BALB/c mice seven days after being immunized subcutaneously twice at 14 day intervals with 100 μg rSsEno emulsified with Freund’s adjuvant) or sera from cats with confirmed sporotrichosis (n=34) obtained from INI/Fundação Oswaldo Cruz, Rio de Janeiro, Brazil. One strip containing Ss16345CWP was incubated with anti-rSsEno. Sera from naïve mice or sera from cats with no evidence of sporotrichosis (n=3) were utilized as negative controls. All sera were diluted 1:100 in PBS. After three washes with PBS, the strips were further incubated for 2 h with goat anti-mouse IgG (Sigma-Aldrich) diluted 1:500 or goat anti-feline IgG (Southern Biotech) diluted 1:1000. Both antibodies were conjugated with horseradish peroxidase (HRP). Protein signals were *visualized* by adding 3,3′-diaminobenzidine *plus* hydrogen peroxide.

### Alignment of enolase sequences

We compared conservation (similarity) between the enolase of *S. schenckii* and the cat and human enolase. The enolase amino acid sequences of *S. schenckii* (GenBank Accession No. ERS97971.1 and *Felis catus* (UniProt Accession: M3 WCP0_FELCA, Homo sapiens (UniProt Accession: P06733) were aligned through the default settings within Clustal Omega^55^.

### Flow cytometry

To demonstrate the enolase on the cell wall of Ss16345, Ss 1099-18, Ss250 and Ss256, 10^6^ yeasts were incubated for 1 h at 37°C with anti-rSsEno serum. Serum from naïve mice was used as a nonspecific binding control at a 1:50 dilution. After incubation, cells were washed twice with PBS for 1 h at 37°C and then incubated with a FITC-conjugated rabbit anti-mouse IgG antibody (Sigma-Aldrich) at a 1:500 dilution. After washing, samples were acquired with the BD Accuri C6 flow cytometer (BD Biosciences). The acquisition threshold was set to 50,000 on FSC-H for debris exclusion, and at least 50,000 events were effectively included in each analysis. Binding of serum antibodies to the yeast cell surface was assessed through the median fluorescence intensity (MFI) on the FL1 channel using the flow cytometer’s proprietary software.

### Electron microscopy

To visualize enolase on the Ss16345, Ss1099-18, Ss250 and Ss256 cell surface, we performed preembedding immunogold experiments using intact yeast cells this fungus, as described previously^43^. Briefly, the yeast cells were fixed with 2.5 glutaraldehyde v/v in 0.1 M cacodylate buffer, pH 7.2, for 24 h at 4°C. Ultrathin sections of each fungus were prepared and treated overnight with the primary antibody (polyclonal anti-rSsEno) diluted 1:100 in PBS at 4°C. The grids were then incubated overnight with the labeled Au-conjugated secondary antibody rabbit IgG (10 nm average particle size, 1:20) at 4°C. The grids were stained with 4% uranyl acetate and lead citrate and observed with a Jeol 1011 transmission electron microscope (Jeol, Tokyo, Japan). Controls were obtained by incubating the ultrathin sections with NS.

### Immunization schedule

BALB/c mice (n = 5) were injected subcutaneously (s.c.) three times in the back of the neck, with 2-week intervening period, with one of the following formulations diluted in 100 μl of PBS: rSsEno100 alone (100 μg), PGA+rSsEno100 [10% Montanide™ Pet Gel A (PGA), SEPPIC, France plus 100 μgrSsEno] or PBS alone as a negative control. One week after the later immunization, mice were euthanized in a CO2 chamber and bled by heart puncture to obtain serum, which was aliquoted and stored at −20°C until use.

### Quantification of the rSsEno-specific antibody response by ELISA

rSsEno IgG, IgG1, IgG2a and IgG3 antibody titration was conducted as described by Portuondo *et al*.^19^ with some modifications. Briefly, a 96-well ELISA plate (Costar) was coated with 5 μg rSsEno/mL in PBS and incubated overnight at 4 °C. The plate was washed with washing buffer (0.1% Tween 20 in PBS) and then saturated for 1 h at RT with blocking buffer (5% dried skim milk in washing buffer). Next, dilutions (1:100 in blocking buffer) of the serum samples were added to each well and incubated for 2 h at RT. After washing, 100 μl/well of peroxidase-conjugated anti-mouse IgG (1/500) (Sigma) in blocking buffer was added and incubated at 37°C for 1 h. For determination of the IgG1, IgG2a and IgG3 subclasses, ELISA plates coated as before were first incubated with an unconjugated rabbit anti-mouse IgG1, IgG2a or IgG3 (Bio-Rad) at 37°C for 1 h and then with a peroxidase-conjugated goat anti-rabbit IgG (Sigma) overnight at 4°C. After exhaustive washing, immune complexes were revealed by incubation with tetramethylbenzidine for 30 min at RT. The reaction was stopped by the addition of 50 μL/well 1 M H_2_SO_4_, and the absorbance was read with an ELISA reader (Multiskan ascent, Labsystem) at 450 nm.

### Cytokine production

To evaluate the cytokine production induced by rSsEno-stimulated spleen cells, splenocytes isolated from each group of animals were harvested seven days after the third immunization. Collected cells were washed, suspended in complete RPMI-1640 medium (cRPMI; RPMI-1640 medium containing 0.02 mM 2-mercaptoethanol, 100 U/mL penicillin, 100 U/mL streptomycin, 2 mM l-glutamine, and 5% fetal bovine serum) and then plated in triplicate in 96-well plates (Costar, USA) to final concentration 2.5×10^6^ cells/mL with 20 μg of rSsEno/mL in cRPMI for 24 h at 37°C with 5% CO_2_. Concanavalin A (0.25 l g/ml) or cRPMI alone were used as positive and negative controls, respectively. Supernatant-accumulated cytokine concentrations (IL-2, IL-10, IL-4, IL-6, IFN-γ, TNF-α, and IL-17A) were simultaneously measured using the mouse Th1/Th2/Th17 cytokine cytometric bead array (CBA) kit (BD Biosciences). Briefly, 50 μL of each standard or supernatant sample was incubated for 2 h at RT with an equal volume of PhycoErythrin (detection reagent) and the mixed capture beads. After incubation, the samples were centrifuged at 200 × g for 5 min, and the pellet was resuspended in 300 μL of wash buffer and analyzed using a flow cytometer (BD Accuri C6, BD Biosciences).

### Protection assay

BALB/c mice (n=10) were immunized according to the immunization schedule described previously. Seven days after the final boost, mice were challenged intravenously with 10^5^ of the highly virulent *S. brasiliensis* Ss250 yeast in 0.1 mL of PBS via the tail vein, as described by Ishida et al.^56^. Animals were monitored daily for 45 days postinfection to determine the survival curve and efficacy of each vaccine formulation.

### Statistical analysis

Data were analyzed using one-way analysis of variance (ANOVA) followed by Tukey’s post-test using Graph Pad Prism 5. In this study, a p value of < 0.05 was considered significant. The results are expressed as the mean ± SD.

## Acknowledgments

This work was supported by Fundação de Amparo à Pesquisa do Estado de São Paulo (FAPESP, grant number 2015/09340-4, 2017/13228-0 and 2017/26774-3). We are also grateful to Dr. Sandro Antonio Pereira (Instituto Nacional de Infectologia Evandro Chagas – INI, Fiocruz, Rio de Janeiro, RJ) for providing the *S. brasiliensis* isolate and sera from cats used in this study.

## Contributions

Conception of the work: D.L.P.F., A.B.D., I.Z.C. Design of research: D.L.P.F., A.B.D., I.Z.C., P.R.D.S; L.S.F., D.T.M Performed experiments: D.L.P.F., A.B.D., P.R.D.S; L.S.F., D.T.M., M.L.A.L.,C.M.M., Data analysis: D.L.P.F., A.B.D., I.Z.C., P.R.D.S; L.S.F., D.T.M., C.M.M.,F.G.Q., J.C.B. Interpreted results of experiments: D.L.P.F., A.B.D., I.Z.C., Prepared figures: D.L.P.F., A.B.D., I.Z.C., Wrote the manuscript: D.L.P.F., A.B.D., I.Z.C. All authors contributed to the final version of the paper and gave final approval for publication.

